# Glial Contribution to the Pathogenesis of Post-Operative Delirium Revealed by Multi-omic Analysis of Brain Tissue from Neurosurgery Patients

**DOI:** 10.1101/2025.03.13.643155

**Authors:** Takaya Ishii, Tao Wang, Kazuki Shibata, Shota Nishitani, Takehiko Yamanashi, Nadia E. Wahba, Tomoteru Seki, Kaitlyn J. Thompson, Kyosuke Yamanishi, Tsuyoshi Nishiguchi, Akiyoshi Shimura, Bun Aoyama, Nipun Gorantla, Nathan J. Phuong, Hieu D. Nguyen, Therese A. Santiago, Yoshitaka Nishizawa, Takaaki Nagao, Mathew A Howard, Hiroto Kawasaki, Kyosuke Hino, Atsushi Ikeda, Michael P. Snyder, Gen Shinozaki

## Abstract

Post-operative delirium (POD) is a common complication after surgery especially in elderly patients, characterized by acute disturbances in consciousness and cognition, which negatively impacts long-term outcomes. Effective treatments remain elusive due to the unclear pathophysiology of POD. To address the knowledge gap, we investigated DNA methylation profiles and gene expression changes in brain cells from POD and non-POD patients who underwent brain resection surgery for medication refractory epilepsy. DNA methylation analysis revealed alteration in epigenetic status of immune and inflammation-related genes. Single-nucleus RNA sequencing (snRNAseq) identified POD-specific glial cell alterations, particularly in microglia, where neuroinflammation was strongly enhanced, consistent with epigenetic findings. Astrocytes exhibited changes in synapse-related functions and migration. Furthermore, downstream analysis indicated similarities between POD-associated glial cell states and pathologies such as encephalitis and dementia. Overall, this study—the first multi-omics analysis of brain tissue from POD patients—provides direct evidence of glial cell contributions to POD pathogenesis, and highlights potential therapeutic targets.

## Introduction

Delirium is a severe neuropsychiatric syndrome characterized by acute impairments in attention and cognition. Among its various subtypes, Post-Operative Delirium (POD) is particularly prevalent in elderly patients following surgery and is associated with increased mortality, prolonged hospitalization, and long-term cognitive decline^1–3^. Despite its clinical significance, the pathophysiological mechanism of POD remains largely unknown, hindering the development of effective preventive and therapeutic strategies. Identifying reliable biomarkers for POD is crucial for improving patient outcomes and also advancing our understanding of its underlying mechanisms.

Epigenetic modifications, particularly DNA methylation changes, have emerged as potential biomarkers for delirium. Our previous studies demonstrated consistent patterns of alterations in DNA methylation levels in blood cells among delirium patients. This was replicated across four independent cohorts^4^. However, the extent to which these changes in peripheral tissues correlate with molecular alterations in the brain remains unclear. Thus, a direct investigation of brain tissue is essential for elucidating the neurobiological mechanisms of POD. In a previous study, we compared DNA methylation profiles across brain, saliva, and blood samples from medication-refractory epilepsy patients undergoing brain resection surgery, stratified by the presence or absence of POD ^5^. While inflammatory signatures potentially linked to delirium were detected in both the brain and blood, their molecular profiles showed limited overlap, underscoring the need for a more detailed analysis of brain tissue. However, studies examining delirium-specific changes in the brain remain scarce due to the difficulty of obtaining brain samples, leaving fundamental aspects of its pathogenesis unexplored.

Single-cell and single-nucleus RNA sequencing (scRNAseq/snRNAseq) provide powerful tools for characterizing the transcriptomic landscape of individual brain cells. These techniques have been widely applied in neurodegenerative and psychiatric disorders^6,7^, offering new insights into pathophysiology, biomarker discovery, and therapeutic targets. While a recent study identified potential POD biomarkers in blood cells using scRNAseq^8^, no research to date has employed this approach to analyze brain tissue in POD patients.

In this study, we employ a multi-omics approach (genome-wide DNA methylation analysis and snRNAseq) to identify delirium-specific molecular signatures in brain tissue. By analyzing brain samples, we aim to uncover the epigenetic and transcriptomic features unique to POD, offering new mechanistic insights into its pathophysiology. This study represents the first application of snRNAseq to human brain tissue in POD, providing a crucial step toward identifying the molecular underpinnings of its pathophysiological mechanisms, leading to the potential therapeutic targets for this serious condition.

## Materials and Methods

### Subject

DNA methylation data and samples derived from a previous cohort study were analyzed ^5,9^. In this cohort, subjects with medication-refractory epilepsy scheduled for brain resection surgery were recruited at the University of Iowa Hospitals and Clinics. This cohort study was approved by the University of Iowa’s Human Subjects Research Institutional Review Board (#200112047 and #20190791). The present study was also approved by the Stanford University Institutional Review Board (#62033). All patients underwent a series of two neurosurgical procedures: electrode placement for the identification of seizure focus and subsequent tissue resection. To identify POD cases, detailed chart reviews of medical records including physical, neurological, and mental status as well as nursing reports were conducted. Patients with post-operative fluctuations in awareness and orientation were considered as positive cases of POD. A board-certified consultation liaison psychiatrist (G.S.) reviewed the cases in question for the final decision on delirium categorization^5,10^.

For this study, 18 cases were selected for analysis and divided into the POD group and the non-POD group. The 9 cases that developed POD after the brain resection surgery were considered the POD group, and the 9 cases that did not develop POD after both surgeries were considered the non-POD group. **Table 1** describes the detailed characteristics of the subjects.

**Table 1.** Subject characteristics.

| Classification | All Subjects | POD | non-POD | *P* | Statistical test |
| --- | --- | --- | --- | --- | --- |
| *N* | 18 | 9 | 9 |  |  |
| Mean age (years) | 36.4 | 39.9 | 30.8 | 0.33 | *t* = 1.007 |
| SD | 14.1 | 16.0 | 11.7 |  |  |
| Sex (Female) (n) | 7 | 4 | 3 | 0.63 | $\chi^2 = 0.2338$ |
| % | 38.9 | 44.4 | 33.3 |  |  |
| BMI (kg/m<sup>2</sup>) | 29.0 | 28.3 | 29.6 | 0.75 | *t* = -0.3251 |
| SD | 8.5 | 7.7 | 9.6 |  |  |
| Mean anesthesia time (min) | 542.6 | 545.9 | 539.2 | 0.84 | *t* = 0.2126<br>(unequal variance) |
| SD | 64.6 | 84.5 | 41.4 |  |  |
| Mean procedure time (min) | 352.2 | 360.0 | 344.4 | 0.68 | *t* = 0.4192<br>(unequal variance) |
| SD | 76.8 | 100.0 | 49.0 |  |  |
| Mean blood loss (mL) | 182.5 | 175.0 | 209.4 | 0.47 | *t* = -0.7436 |
| SD | 151.7 | 171.7 | 133.3 |  |  |
| History of alcohol use (n) | 6 | 3 | 3 | 1.00 | Fisher's exact test<br>OR= 1.000, 95% CI<br>[0.0919,10.88] |
| % | 33.3 | 33.3 | 33.3 |  |  |
| History of tobacco use (n) | 8 | 5 | 3 | 0.64 | Fisher's exact test<br>OR= 2.372, 95% CI<br>[0.2652, 25.53] |
| % | 44.4 | 55.5 | 33.3 |  |  |

BMI, Body mass index; POD, post-operative delirium; SD, Standard deviation; OR, Odds Ratio; CI, Confidence Interval

### Brain sample collection and processing

Brain samples were immediately stored at −80℃, i.e., flash-freezing, after brain resection. For snRNAseq analysis, nuclei were isolated from the frozen tissues as previously described ^11^. Briefly, frozen tissue was homogenized in a chilled lysis buffer using a Dounce homogenizer, followed by differential centrifugation with iodixanol gradient separation to isolate the nuclei. The nuclei pellet was then carefully collected, resuspended, and subjected to additional washes and centrifugation steps. The final nuclei were resuspended in PBS/1% BSA, and utilized for further analysis. For DNA methylation analysis, genomic DNA was extracted from bulk samples using the MasterPure DNA extraction kit (Epicentre) as described in previous publications ^5^.

### DNA methylation analysis

DNA derived from brain tissues was bisulfite-converted with the EZ DNA Methylation Kit (Zymo Research). Genome-wide DNA methylation was analyzed using the Infinium HumanMethylationEPIC BeadChip Kit (Illumina). The arrays were scanned with the Illumina iScan platform. Raw methylation data was processed on R software^12^ using the R packages “ChAMP”^13,14^ and “Minfi”^15,16^. Probes were excluded if (i) the detection *p*-value exceeded 0.01, (ii) fewer than three beads were present in at least 5% of samples per probe, (iii) the probes were non-CpG, SNP-related, or multi-hit, or (iv) the probes were located on the X or Y chromosomes. Normalization was performed using beta-mixture quantile dilation^17^. To remove chip batch effects, we used champ.runCombat with protecting group from the removal process. The R package “Rnbeads”^18^ employing the limma method^19^ was used to perform differential methylation analysis. Age, sex, time under anesthesia, blood loss, smoking status, and alcohol use were included as covariates in the analysis.

### snRNAseq sequencing library preparation

10,000 nuclei were applied for library preparation using Chromium Next GEM Single Cell Multiome Reagent Kit (10x Genomics) following the manufacturer’s protocol. The libraries were pooled and sequenced in the Illumina NovaSeq PE150 system targeting an average of 50,000 reads per nucleus.

### snRNAseq data processing and integration

The raw reads were mapped to human reference genome version GRCh38 and unique molecular identifiers (UMIs) counts for each gene in each cell by CellRanger Arch v2.0.0. (10x Genomics). We used the R(v4.3.2) package Seurat (v.5.0.2)^20^ for processing the raw count matrices. Only genes expressed in at least 3 nuclei were considered. For each sample, nuclei were excluded if ( i) presented unique genes were fewer than 200 or more than 6,000, or ( ii) total UMI counts were less than 500 or over 25,000, or (iii) mitochondrial RNA content was superior to 5%. The individual Seurat objects for each sample were lognormalized using the Seurat function “NormalizeData” with 2,000 variable genes. Then, all samples were integrated by the following functions, “SelectIntegrationFeatures”, “PrepSCTIntegration”, “FindIntegrationAnchors” and “IntegrateData”.

### snRNAseq data analysis

Integrated data were utilized for downstream analysis. Age, gender, and mitochondrial content were regressed during the data scaling process. Dimensionality reduction was performed using Principal Component Analysis (PCA) and Uniform Manifold Approximation and Projection (UMAP), incorporating the top 20 principal components (PCs). Cell clusters were identified using the “FindClusters” function. Each cluster was annotated with cell types based on representative marker genes corresponding to major brain cell types, including microglia, astrocytes, oligodendrocytes, oligodendrocyte progenitor cells (OPCs), endothelial cells, excitatory neurons, and inhibitory neurons ^21–23^. Clusters expressing markers of multiple cell types were identified as multiplets and excluded. For cell type-specific analysis, clusters corresponding to each cell type were independently extracted and subjected to scaling, dimensional reduction, and clustering in the same manner. Differential gene expression (DEG) analysis was performed with the “Findmarkers” function. In this study, genes meeting the following criteria were defined as DEGs: adjusted p-value < 0.05, Log2 Fold Change > 0.25, and pct > 0.1 (That means that the gene is expressed in more than 10% of cells in at least one of the two groups (POD and non-POD group) being compared). Trajectory analysis was performed using the “monocle3” package ^24–27^.

### Gene Ontology (GO) and pathway analysis

GO analysis for enriched genes in snRNAseq analysis was performed using Metascape (v3.5 20240901) ^28^, and pathway analysis was performed with QIAGEN Ingenuity Pathway Analysis (IPA) (Version 127006219, Build: ing_ruby). As for DNA methylation data analysis, R package missMethyl ^29^ was used for GO and pathway analysis by adjusting for the variable number of CpG sites tested in each gene.

### Statistical analysis

Most statistical analyses were performed using R software. Regarding enrichment analysis for snRNAseq, p-values were calculated by each platform (Metascape and IPA).

## Results

### Participant demographics

A total of 18 patients (9 POD, 9 non-POD) who underwent brain resection neurosurgery were analyzed. There was no significant difference between the POD and the non-POD groups in each baseline characteristic **(Table 1)**. Brain samples were taken from various brain regions, including the temporal cortex, frontal cortex, hippocampus, amygdala, and occipital lobe depending on the location of seizure focus.

### DNA methylation analysis

DNA methylation analysis at 700,284 CpG sites was determined using the EPIC Infinium Bead as depicted in **Figure 1A**. DNA methylation status was compared between the POD and non-POD groups and ranked by significance based on ascending p-values **(Table S1)**. Even the most significantly different site; cg18199231[*GTF2IRD1*] did not reach statistical significance after p-value adjustment correcting for genome-wide multiple comparison. Given the absence of genome-wide significant signals, we proceeded with further analysis, focusing on CpGs with relatively large changes. In this analysis, differentially methylated CpGs (DMCs) were defined based on criteria referenced from the previous study^30^, as CpGs with a p-value < 0.05 and a methylation difference (delta beta-value) > 4% between the two groups. 3,774 DMCs were identified **(Figure 1B)**. PCA plot **(Figure 1C)** and hierarchical clustering heatmap **(Figure 1D)** based on the DMCs demonstrated a clear separation between most POD and non-POD subjects. While the CpG context and gene region distribution of DMCs resembled that of all CpGs **(Figure S1 A, B)**, their methylation rate distribution differed **(Figure 1E)**. Notably, the changes in DNA methylation levels between the non-POD and POD groups varied depending on the initial DNA methylation levels in the non-POD group **(Figure 1F, G)**. Most low-methylated DMCs showed increased DNA methylation in the POD group, whereas high-methylated DMCs displayed both hypermethylation and hypomethylation, with demethylation being more common. Furthermore, high-methylated DMCs contained fewer island-related regions **(Figure S1C)** and transcription start site (TSS) neighbors **(Fig S1E)**, whereas low-methylated DMCs were more frequently associated with island-related regions **(Fig S1D)** and TSS neighbors **(Fig S1F)**.

**Figure 1.**
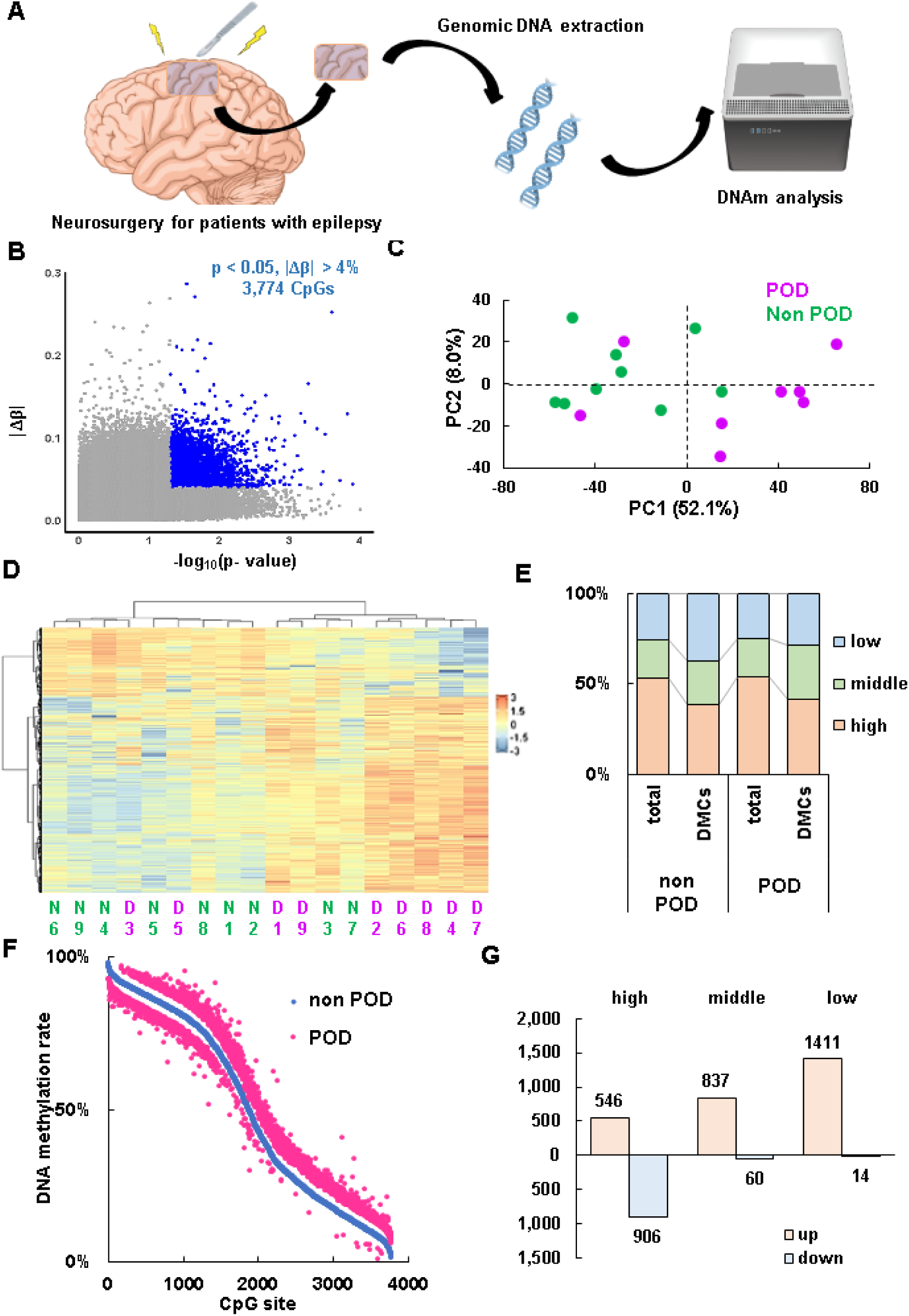
DNA methylation changes between non POD and POD grou. **A** Research scheme for DNA methylation analysis using resected brain tissu **B** Volcano plot displaying 3,774 differently methylated CpGs (DMCs) with a p-value < 0.05 and delta beta value > 4 **C** Principal component analysis of DMCs using Z-score of the beta value **D** Hierarchical clustering heatmap representing DNA methylation of DMCs. Z-score of the beta values is shown. N1-9 represent non-POD subjects and D1-9 represent POD subjects. **E** Distribution of methylation rate; high: 70%>, low: 30%<. Middle: 30-70% **F** The plot shows the methylation rates (β-values) of CpG sites in the POD group (blue) and non-POD group (magenta). The x-axis represents individual CpG sites, while the y-axis indicates the methylation rate. **G** DNA methylation rate of each DMC

To assess whether the observed methylation pattern differences contribute to variations in associated gene groups, we performed Gene Ontology (GO) and pathway analyses. Although no GO terms reached statistical significance after FDR correction, several immune-related terms, including T cell differentiation and T cell activation, were ranked among the top (p < 0.01), alongside terms related to cell adhesion. Similarly, pathway analysis identified immune-related pathways, such as the chemokine signaling pathway and human cytomegalovirus infection, as top-ranking **(Table 2)**. Since aging is a major risk factor for POD, we further examined the methylation status of age-associated CpGs. In a recent large-scale study analyzing over 10,000 human samples (manuscript in preparation), we identified 24,744 CpGs with genome-wide significant (p < 5.78 × 10⁻⁸) and strong (R² > 0.5) age-associated methylation changes. Among these, 152 overlapped with the identified DMCs in the present study. Although no GO or pathway terms reached FDR significance in the analysis of *age-associated DMCs*, immune-and inflammation-related terms were among the top-ranked **(Table 3)**. These findings suggest that neuroinflammatory alterations are a key feature of the POD group.

**Table 2.**
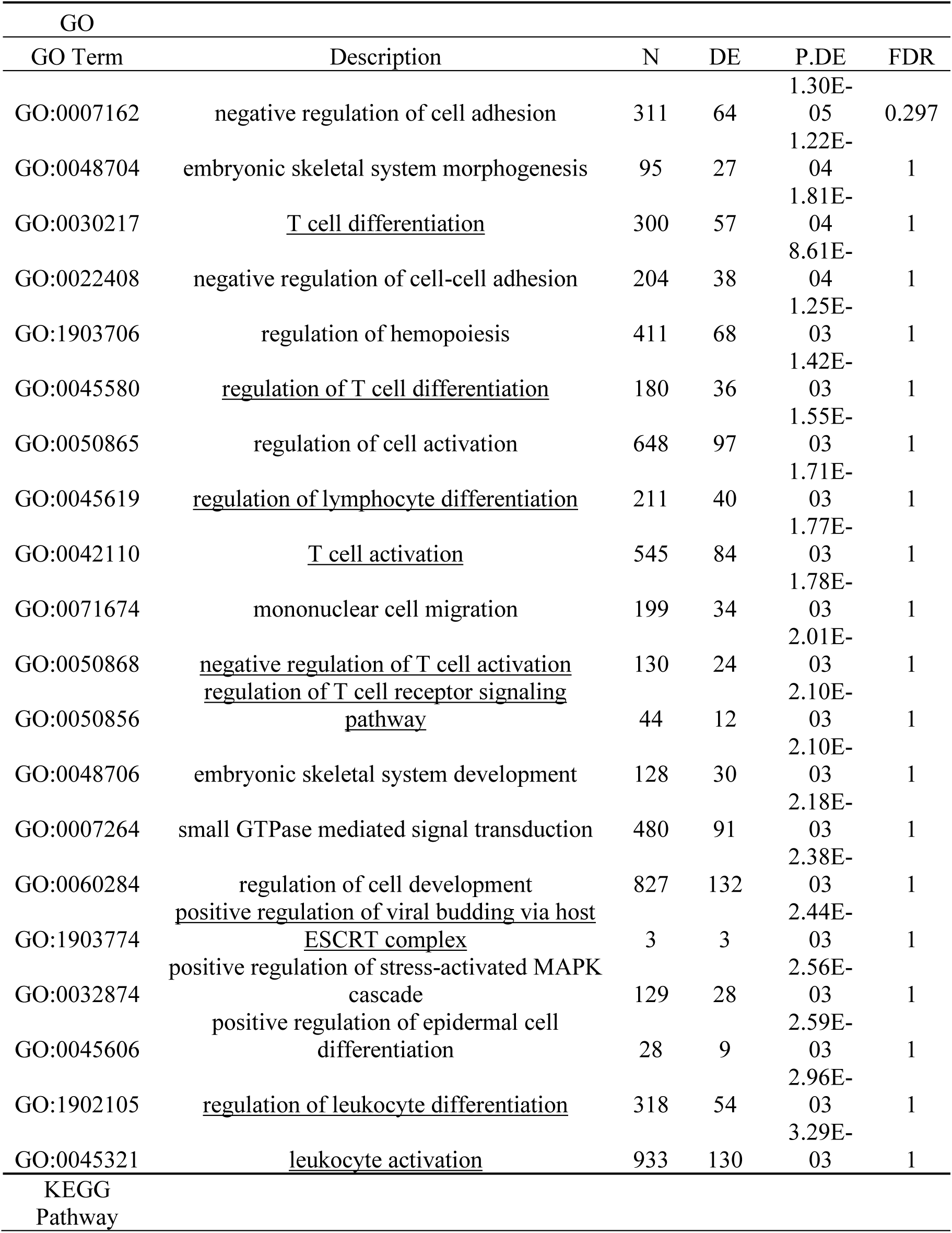
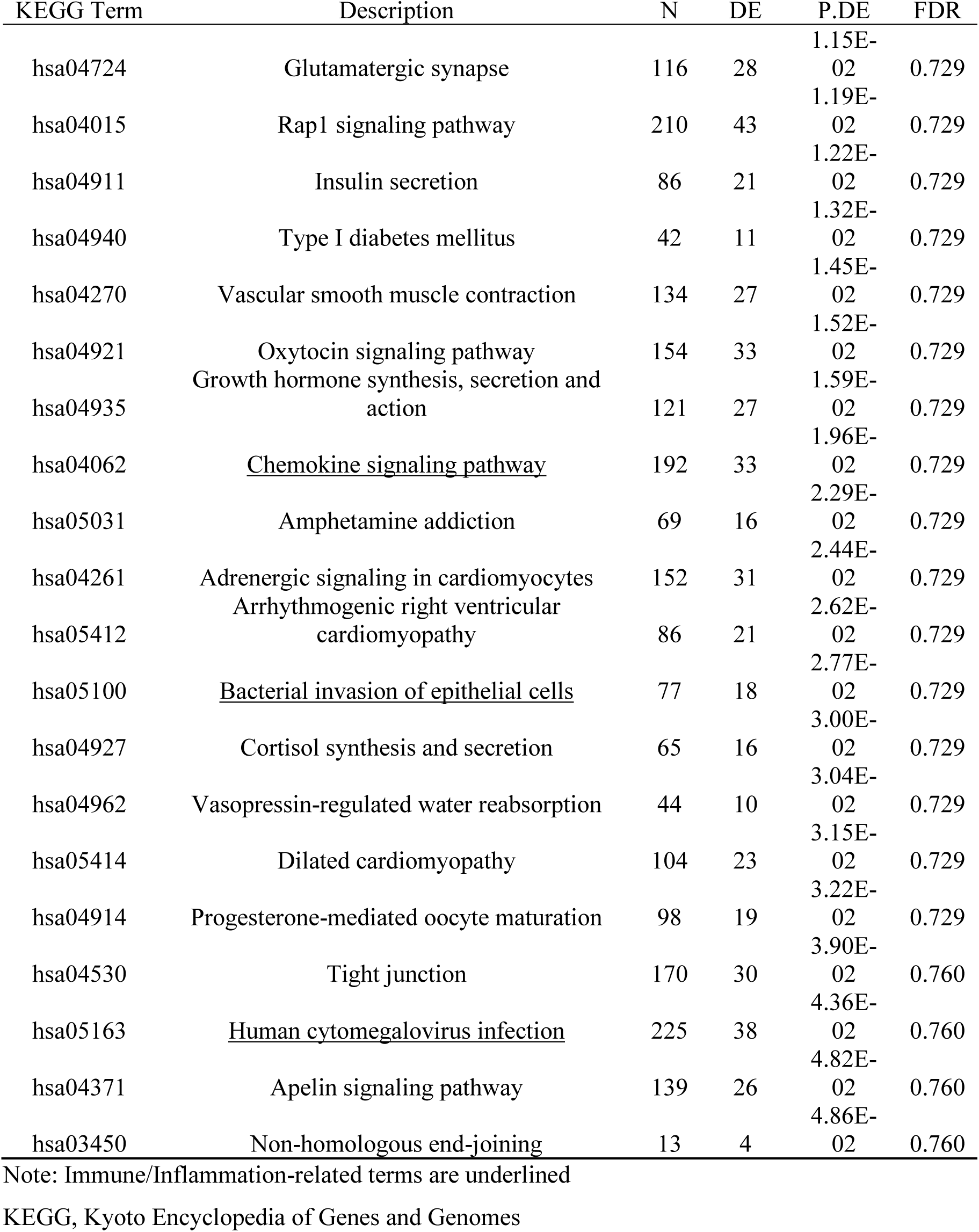
Top 20 terms of GO BP and KEGG pathway analysis with all differentially methylated CpGs (p <0.05)

| GO |  |  |  |  |  |
| --- | --- | --- | --- | --- | --- |
| GO Term | Description | N | DE | P.DE | FDR |
| GO:0007162 | negative regulation of cell adhesion | 311 | 64 | 1.30E-05 | 0.297 |
| GO:0048704 | embryonic skeletal system morphogenesis | 95 | 27 | 1.22E-04 | 1 |
| GO:0030217 | <u>T cell differentiation</u> | 300 | 57 | 1.81E-04 | 1 |
| GO:0022408 | negative regulation of cell-cell adhesion | 204 | 38 | 8.61E-04 | 1 |
| GO:1903706 | regulation of hemopoiesis | 411 | 68 | 1.25E-03 | 1 |
| GO:0045580 | <u>regulation of T cell differentiation</u> | 180 | 36 | 1.42E-03 | 1 |
| GO:0050865 | regulation of cell activation | 648 | 97 | 1.55E-03 | 1 |
| GO:0045619 | <u>regulation of lymphocyte differentiation</u> | 211 | 40 | 1.71E-03 | 1 |
| GO:0042110 | <u>T cell activation</u> | 545 | 84 | 1.77E-03 | 1 |
| GO:0071674 | mononuclear cell migration | 199 | 34 | 1.78E-03 | 1 |
| GO:0050868 | <u>negative regulation of T cell activation</u> | 130 | 24 | 2.01E-03 | 1 |
| GO:0050856 | <u>regulation of T cell receptor signaling pathway</u> | 44 | 12 | 2.10E-03 | 1 |
| GO:0048706 | embryonic skeletal system development | 128 | 30 | 2.10E-03 | 1 |
| GO:0007264 | small GTPase mediated signal transduction | 480 | 91 | 2.18E-03 | 1 |
| GO:0060284 | regulation of cell development | 827 | 132 | 2.38E-03 | 1 |
| GO:1903774 | <u>positive regulation of viral budding via host ESCRT complex</u> | 3 | 3 | 2.44E-03 | 1 |
| GO:0032874 | positive regulation of stress-activated MAPK cascade | 129 | 28 | 2.56E-03 | 1 |
| GO:0045606 | positive regulation of epidermal cell differentiation | 28 | 9 | 2.59E-03 | 1 |
| GO:1902105 | <u>regulation of leukocyte differentiation</u> | 318 | 54 | 2.96E-03 | 1 |
| GO:0045321 | <u>leukocyte activation</u> | 933 | 130 | 3.29E-03 | 1 |
| KEGG Pathway |  |  |  |  |  |

| KEGG Term | Description | N | DE | P.DE | FDR |
| --- | --- | --- | --- | --- | --- |
| hsa04724 | Glutamatergic synapse | 116 | 28 | 1.15E-02 | 0.729 |
| hsa04015 | Rap1 signaling pathway | 210 | 43 | 1.19E-02 | 0.729 |
| hsa04911 | Insulin secretion | 86 | 21 | 1.22E-02 | 0.729 |
| hsa04940 | Type I diabetes mellitus | 42 | 11 | 1.32E-02 | 0.729 |
| hsa04270 | Vascular smooth muscle contraction | 134 | 27 | 1.45E-02 | 0.729 |
| hsa04921 | Oxytocin signaling pathway | 154 | 33 | 1.52E-02 | 0.729 |
| hsa04935 | Growth hormone synthesis, secretion and action | 121 | 27 | 1.59E-02 | 0.729 |
| hsa04062 | <u>Chemokine signaling pathway</u> | 192 | 33 | 1.96E-02 | 0.729 |
| hsa05031 | Amphetamine addiction | 69 | 16 | 2.29E-02 | 0.729 |
| hsa04261 | Adrenergic signaling in cardiomyocytes | 152 | 31 | 2.44E-02 | 0.729 |
| hsa05412 | Arrhythmogenic right ventricular cardiomyopathy | 86 | 21 | 2.62E-02 | 0.729 |
| hsa05100 | <u>Bacterial invasion of epithelial cells</u> | 77 | 18 | 2.77E-02 | 0.729 |
| hsa04927 | Cortisol synthesis and secretion | 65 | 16 | 3.00E-02 | 0.729 |
| hsa04962 | Vasopressin-regulated water reabsorption | 44 | 10 | 3.04E-02 | 0.729 |
| hsa05414 | Dilated cardiomyopathy | 104 | 23 | 3.15E-02 | 0.729 |
| hsa04914 | Progesterone-mediated oocyte maturation | 98 | 19 | 3.22E-02 | 0.729 |
| hsa04530 | Tight junction | 170 | 30 | 3.90E-02 | 0.760 |
| hsa05163 | <u>Human cytomegalovirus infection</u> | 225 | 38 | 4.36E-02 | 0.760 |
| hsa04371 | Apelin signaling pathway | 139 | 26 | 4.82E-02 | 0.760 |
| hsa03450 | Non-homologous end-joining | 13 | 4 | 4.86E-02 | 0.760 |

Note: Immune/Inflammation-related terms are underlined
KEGG, Kyoto Encyclopedia of Genes and Genomes

**Table 3.** Top terms of GO BP and KEGG analysis with differentially methylated CpGs (age-associated CpGs)

| GO |  |  |  |  |  |
| --- | --- | --- | --- | --- | --- |
| GO Term | Description | N | DE | P.DE | FDR |
| GO:0002876 | <u>positive regulation of chronic inflammatory response to antigenic stimulus</u> | 2 | 2 | 1.14E-05 | 0.194 |
| GO:0002439 | <u>chronic inflammatory response to antigenic stimulus</u> | 3 | 2 | 2.55E-05 | 0.194 |
| GO:0002874 | <u>regulation of chronic inflammatory response to antigenic stimulus</u> | 3 | 2 | 2.55E-05 | 0.194 |
| GO:0002678 | <u>positive regulation of chronic inflammatory response</u> | 4 | 2 | 6.52E-05 | 0.373 |
| GO:0032741 | <u>positive regulation of interleukin-18 production</u> | 7 | 2 | 1.86E-04 | 0.853 |
| GO:1902565 | <u>positive regulation of neutrophil activation</u> | 6 | 2 | 3.04E-04 | 1 |
| GO:0002925 | <u>positive regulation of humoral immune response mediated by circulating immunoglobulin</u> | 7 | 2 | 3.93E-04 | 1 |
| GO:2000343 | <u>positive regulation of chemokine (C-X-C motif) ligand 2 production</u> | 9 | 2 | 6.60E-04 | 1 |
| GO:0032621 | <u>interleukin-18 production</u> | 13 | 2 | 6.76E-04 | 1 |
| GO:0032661 | <u>regulation of interleukin-18 production</u> | 13 | 2 | 6.76E-04 | 1 |
| GO:1903223 | positive regulation of oxidative stress-induced neuron death | 8 | 2 | 6.87E-04 | 1 |
| GO:0002676 | <u>regulation of chronic inflammatory response</u> | 10 | 2 | 7.60E-04 | 1 |
| GO:0032755 | <u>positive regulation of interleukin-6 production</u> | 99 | 4 | 9.00E-04 | 1 |
| GO:0071803 | positive regulation of podosome assembly | 10 | 2 | 1.10E-03 | 1 |
| GO:0070305 | response to cGMP | 9 | 2 | 1.71E-03 | 1 |
| GO:0060251 | regulation of glial cell proliferation | 39 | 3 | 1.84E-03 | 1 |
| GO:0002863 | <u>positive regulation of inflammatory response to antigenic stimulus</u> | 13 | 2 | 1.95E-03 | 1 |
| GO:0033477 | S-methylmethionine metabolic process | 1 | 1 | 1.97E-03 | 1 |
| GO:1902563 | regulation of neutrophil activation | 15 | 2 | 2.03E-03 | 1 |
| GO:0042756 | drinking behavior | 12 | 2 | 2.07E-03 | 1 |
| KEGG Pathway |  |  |  |  |  |

| KEGG Term | Description | N | DE | P.DE | FDR |
| --- | --- | --- | --- | --- | --- |
| hsa04940 | Type I diabetes mellitus | 42 | 3 | 2.13E-03 | 0.617 |
| hsa04625 | <u>C-type lectin receptor signaling pathway</u> | 104 | 4 | 3.41E-03 | 0.617 |
| hsa04144 | Endocytosis | 249 | 6 | 5.60E-03 | 0.676 |
| hsa05143 | African trypanosomiasis | 36 | 2 | 1.15E-02 | 0.799 |
| hsa05152 | <u>Tuberculosis</u> | 178 | 4 | 1.19E-02 | 0.799 |
| hsa05235 | <u>PD-L1 expression and PD-1 checkpoint pathway in cancer</u> | 89 | 3 | 1.75E-02 | 0.799 |
| hsa04064 | <u>NF-kappa B signaling pathway</u> | 104 | 3 | 1.80E-02 | 0.799 |
| hsa05142 | <u>Chagas disease</u> | 102 | 3 | 1.89E-02 | 0.799 |
| hsa05144 | <u>Malaria</u> | 50 | 2 | 1.99E-02 | 0.799 |
| hsa04650 | <u>Natural killer cell mediated cytotoxicity</u> | 125 | 3 | 2.64E-02 | 0.871 |
| hsa04061 | <u>Viral protein interaction with cytokine and cytokine receptor</u> | 99 | 2 | 2.65E-02 | 0.871 |
| hsa05321 | <u>Inflammatory bowel disease</u> | 64 | 2 | 4.28E-02 | 1 |
| hsa04660 | <u>T cell receptor signaling pathway</u> | 118 | 3 | 4.61E-02 | 1 |

Note: Immune/Inflammation-related terms are underlined

To assess whether these trends were influenced by the original methylation rate and its pattern of change, we analyzed DMCs based on their initial methylation status. Among low-methylated DMCs that exhibited increased methylation in the POD group (1,411 CpGs), the top two GO terms were associated with embryonic skeletal system development, which were also enriched in the analysis of all DMCs. Developmental processes were prominently ranked among the top terms. Although no pathways reached statistical significance, several overlapped with those identified in the analysis of all DMCs, including the Rap1 signaling pathway, insulin signaling pathway, and oxytocin signaling pathway **(Table 4)**. In contrast, GO and pathway analyses of highly methylated DMCs that underwent demethylation in the POD group (906 CpGs) revealed a distinct trend. GO analysis showed that most top-ranking biological process terms were related to immunity and inflammation, with “adaptive immune response” approaching statistical significance (FDR = 0.0595). A similar pattern was observed in pathway analysis, where immune- and inflammation-related pathways were predominant, including the NF-kappa B signaling pathway, which reached statistical significance **(Table 5)**. These findings suggest that enhanced demethylation of highly methylated CpG sites in immune- and inflammation-related genes is a hallmark of the POD group, indicating that neuroinflammatory changes in POD may be epigenetically regulated.

**Table 4.** Top 20 terms of GO BP and KEGG analysis with differentially methylated CpGs (high-methylated, down)

| GO |  |  |  |  |  |
| --- | --- | --- | --- | --- | --- |
| GO Term | Description | N | DE | P.DE | FDR |
| GO:0002250 | <u>adaptive immune response</u> | 450 | 28 | 2.60E-06 | 0.059 |
| GO:0032874 | positive regulation of stress-activated MAPK cascade | 129 | 15 | 9.11E-06 | 0.074 |
| GO:0050856 | <u>regulation of T cell receptor signaling pathway</u> | 44 | 8 | 1.13E-05 | 0.074 |
| GO:0070304 | positive regulation of stress-activated protein kinase signaling cascade | 131 | 15 | 1.29E-05 | 0.074 |
| GO:0050854 | regulation of antigen receptor-mediated signaling pathway | 65 | 9 | 4.33E-05 | 0.167 |
| GO:0046330 | <u>positive regulation of JNK cascade</u> | 95 | 12 | 5.86E-05 | 0.167 |
| GO:0042110 | <u>T cell activation</u> | 545 | 31 | 6.55E-05 | 0.167 |
| GO:0046649 | <u>lymphocyte activation</u> | 768 | 39 | 7.85E-05 | 0.180 |
| GO:0045321 | <u>leukocyte activation</u> | 933 | 44 | 9.48E-05 | 0.185 |
| GO:0022408 | negative regulation of cell-cell adhesion | 204 | 16 | 9.71E-05 | 0.185 |
| GO:1903038 | negative regulation of leukocyte cell-cell adhesion | 149 | 12 | 1.44E-04 | 0.236 |
| GO:0002252 | <u>immune effector process</u> | 628 | 30 | 1.75E-04 | 0.264 |
| GO:0050868 | <u>negative regulation of T cell activation</u> | 130 | 11 | 1.85E-04 | 0.264 |
| GO:0032872 | regulation of stress-activated MAPK cascade | 192 | 16 | 2.75E-04 | 0.336 |
| GO:0002876 | <u>positive regulation of chronic inflammatory response to antigenic stimulus</u> | 2 | 2 | 2.77E-04 | 0.336 |
| GO:0098609 | cell-cell adhesion | 945 | 48 | 2.79E-04 | 0.336 |
| GO:1901222 | <u>regulation of NIK/NF-kappaB signaling</u> | 110 | 9 | 3.28E-04 | 0.368 |
| GO:0050860 | <u>negative regulation of T cell receptor signaling pathway</u> | 25 | 5 | 3.37E-04 | 0.368 |
| GO:0070302 | regulation of stress-activated protein kinase signaling cascade | 195 | 16 | 3.56E-04 | 0.371 |
| GO:0006955 | <u>immune response</u> | 1631 | 58 | 3.87E-04 | 0.376 |
| KEGG Pathway |  |  |  |  |  |

| KEGG Term | Description | N | DE | P.DE | FDR |
| --- | --- | --- | --- | --- | --- |
| hsa04064 | <u>NF-kappa B signaling pathway</u> | 104 | 11 | 1.07E-04 | 0.039 |
| hsa04659 | <u>Th17 cell differentiation</u> | 107 | 9 | 6.61E-03 | 0.542 |
| hsa04976 | Bile secretion | 89 | 7 | 8.71E-03 | 0.542 |
| hsa04940 | Type I diabetes mellitus | 42 | 5 | 8.80E-03 | 0.542 |
| hsa04060 | <u>Cytokine-cytokine receptor interaction</u> | 293 | 11 | 8.87E-03 | 0.542 |
| hsa04514 | Cell adhesion molecules | 153 | 11 | 9.89E-03 | 0.542 |
| hsa04061 | <u>Viral protein interaction with cytokine and cytokine receptor</u> | 99 | 5 | 1.05E-02 | 0.542 |
| hsa04270 | Vascular smooth muscle contraction | 134 | 10 | 1.20E-02 | 0.542 |
| hsa04668 | <u>TNF signaling pathway</u> | 118 | 8 | 1.68E-02 | 0.637 |
| hsa04650 | <u>Natural killer cell mediated cytotoxicity</u> | 125 | 8 | 1.76E-02 | 0.637 |
| hsa05321 | <u>Inflammatory bowel disease</u> | 64 | 5 | 2.43E-02 | 0.735 |
| hsa05310 | <u>Asthma</u> | 29 | 3 | 2.53E-02 | 0.735 |
| hsa04914 | Progesterone-mediated oocyte maturation | 98 | 7 | 2.83E-02 | 0.735 |
| hsa04062 | <u>Chemokine signaling pathway</u> | 192 | 11 | 2.84E-02 | 0.735 |
| hsa05152 | <u>Tuberculosis</u> | 178 | 9 | 3.48E-02 | 0.829 |
| hsa04625 | <u>C-type lectin receptor signaling pathway</u> | 104 | 7 | 3.66E-02 | 0.829 |
| hsa04080 | Neuroactive ligand-receptor interaction | 366 | 13 | 4.01E-02 | 0.855 |
| hsa04622 | <u>RIG-I-like receptor signaling pathway</u> | 72 | 4 | 4.96E-02 | 0.990 |

Note: Immune/Inflammation-related terms are underlined.

**Table 5.** Top 20 terms of GO BP and KEGG analysis with differentially methylated CpGs (low-methylated, up)

| GO |  |  |  |  |  |
| --- | --- | --- | --- | --- | --- |
| GO Term | Description | N | DE | P.DE | FDR |
| GO:0048704 | embryonic skeletal system morphogenesis | 95 | 21 | 2.30E-07 | 0.005 |
| GO:0048706 | embryonic skeletal system development | 128 | 24 | 9.39E-07 | 0.011 |
| GO:0048705 | skeletal system morphogenesis | 226 | 30 | 5.34E-05 | 0.262 |
| GO:0009952 | anterior/posterior pattern specification | 209 | 26 | 8.94E-05 | 0.336 |
| GO:0048857 | neural nucleus development | 64 | 12 | 1.17E-04 | 0.336 |
| GO:0007399 | nervous system development | 2492 | 186 | 2.57E-04 | 0.478 |
| GO:0045606 | positive regulation of epidermal cell differentiation | 28 | 7 | 3.24E-04 | 0.484 |
| GO:0010159 | specification of animal organ position | 3 | 3 | 3.35E-04 | 0.484 |
| GO:0003002 | regionalization | 421 | 41 | 3.59E-04 | 0.484 |
| GO:0007389 | pattern specification process | 466 | 45 | 3.64E-04 | 0.484 |
| GO:0048523 | negative regulation of cellular process | 4882 | 291 | 6.03E-04 | 0.640 |
| GO:0032026 | response to magnesium ion | 19 | 6 | 6.90E-04 | 0.640 |
| GO:0048598 | embryonic morphogenesis | 612 | 56 | 7.87E-04 | 0.640 |
| GO:0031327 | negative regulation of cellular biosynthetic process | 1656 | 110 | 8.11E-04 | 0.640 |
| GO:0000122 | negative regulation of transcription by RNA polymerase II | 949 | 73 | 8.70E-04 | 0.641 |
| GO:0061140 | lung secretory cell differentiation | 11 | 4 | 8.96E-04 | 0.641 |
| GO:2000480 | negative regulation of cAMP-dependent protein kinase activity | 13 | 5 | 9.36E-04 | 0.641 |
| GO:0045684 | positive regulation of epidermis development | 33 | 7 | 9.53E-04 | 0.641 |
| GO:0072125 | negative regulation of glomerular mesangial cell proliferation | 4 | 3 | 1.03E-03 | 0.641 |
| GO:0009890 | negative regulation of biosynthetic process | 1686 | 111 | 1.03E-03 | 0.641 |
| KEGG Pathway |  |  |  |  |  |

| KEGG Term | Description | N | DE | P.DE | FDR |
| --- | --- | --- | --- | --- | --- |
| hsa04919 | Thyroid hormone signaling pathway | 121 | 19 | 4.46E-04 | 0.162 |
| hsa04015 | Rap1 signaling pathway | 210 | 24 | 5.28E-03 | 0.538 |
| hsa04935 | Growth hormone synthesis, secretion and action | 121 | 16 | 5.79E-03 | 0.538 |
| hsa04970 | Salivary secretion | 93 | 11 | 9.72E-03 | 0.538 |
| hsa04911 | Insulin secretion | 86 | 12 | 1.06E-02 | 0.538 |
| hsa04927 | Cortisol synthesis and secretion | 65 | 10 | 1.14E-02 | 0.538 |
| hsa04211 | Longevity regulating pathway | 89 | 12 | 1.23E-02 | 0.538 |
| hsa04918 | Thyroid hormone synthesis | 75 | 10 | 1.38E-02 | 0.538 |
| hsa04261 | Adrenergic signaling in cardiomyocytes | 152 | 17 | 1.72E-02 | 0.538 |
| hsa04961 | Endocrine and other factor-regulated calcium reabsorption | 53 | 8 | 2.24E-02 | 0.538 |
| hsa05221 | Acute myeloid leukemia | 67 | 9 | 2.36E-02 | 0.538 |
| hsa03450 | Non-homologous end-joining | 13 | 3 | 2.41E-02 | 0.538 |
| hsa04072 | Phospholipase D signaling pathway | 147 | 17 | 2.42E-02 | 0.538 |
| hsa04928 | Parathyroid hormone synthesis, secretion and action | 115 | 14 | 2.45E-02 | 0.538 |
| hsa04022 | cGMP-PKG signaling pathway | 165 | 17 | 2.51E-02 | 0.538 |
| hsa04921 | Oxytocin signaling pathway | 154 | 17 | 2.53E-02 | 0.538 |
| hsa04962 | Vasopressin-regulated water reabsorption | 44 | 6 | 2.66E-02 | 0.538 |
| hsa04915 | Estrogen signaling pathway | 137 | 14 | 2.68E-02 | 0.538 |
| hsa05205 | Proteoglycans in cancer | 202 | 20 | 2.85E-02 | 0.544 |
| hsa04371 | Apelin signaling pathway | 139 | 14 | 3.75E-02 | 0.600 |

Note: Immune/Inflammation-related terms are underlined

### snRNAseq analysis

Although DNA methylation analysis identified unique characteristics associated with POD, the data represented a mixture of multiple cell types from bulk brain tissues. To investigate POD-related changes at the single-cell level, we performed snRNA-seq analysis using brain tissues from the same subjects. Nuclei were isolated from frozen resected brain tissue of the same 18 subjects, followed by snRNA-seq analysis **(Figure 2A)**. After quality control filtering and clustering (see Methods), we retained 95,017 nuclei, which were categorized into seven distinct cell-type clusters based on the expression of representative marker genes **(Figure 2B, C)**. While the distribution of each cell type varied across samples, all 18 subjects contributed a certain number of nuclei for analysis across all identified cell types and each cell type was present in a consistent proportion in both non-POD and POD groups **(Figure 2D, E)**.

**Figure 2.**
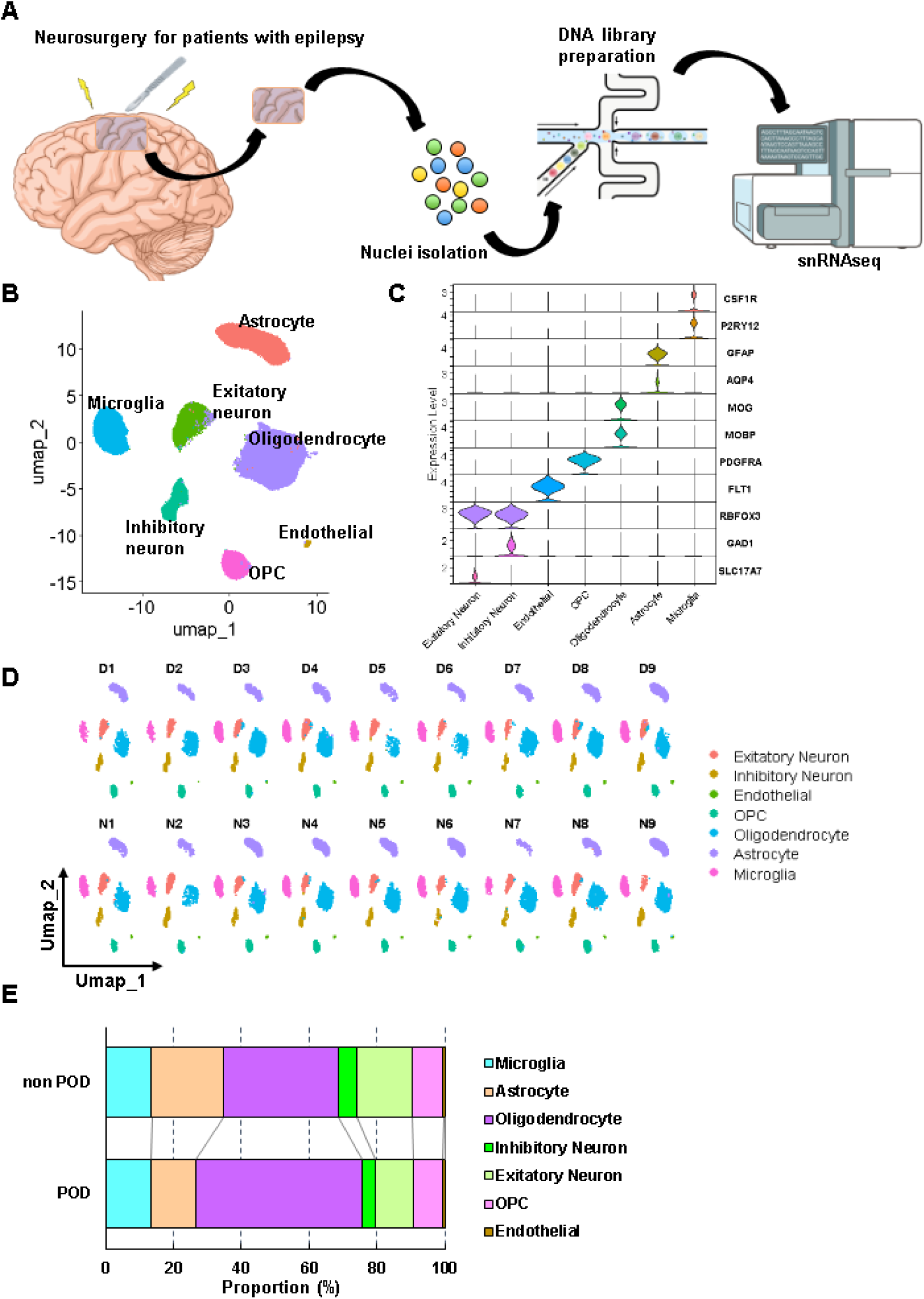
snRNAseq analysis for microglia cluster. **A** Research scheme for snRNAseq analysis using resected brain tissue **B** UMAP for all nuclei colored by cell type annotations. **C** Representative cell type specific marker genes expression **D** UMAPs split by each sample. **E** Proportions of each cell type

### Transcriptional profiling of microglia reveals the upregulation of neuroinflammation in the POD group

Glial cells have been reported to play a crucial role in the pathogenesis of POD through neuroinflammation and other mechanisms^31^, and our DNA methylation analysis suggests that neuroinflammation may be exacerbated in the POD group. Additionally, LPS-injected animal models and surgical models, both of which induce neuroinflammation with elevated cytokines and activated microglia, are widely recognized as classic animal models of delirium ^32,33^. Therefore, we analyzed glial cell clusters from the human brain to investigate the relationship between the transcriptional state of glial cells and POD, with a particular focus on the neuroinflammatory state in the POD group. First, we analyzed microglial clusters, as microglia are the key cell type involved in neuroinflammation. A total of 12,859 nuclei were identified as microglia, characterized by the expression of representative markers CSF1R and P2RY12 **(Figure 3A**, **Figure 2C)**. Compared to the non-POD group, microglia in the POD group exhibited 332 significantly differentially expressed genes (DEGs), with 291 genes upregulated and 41 genes downregulated **(Figure 3B, Table S2)**. Gene Ontology (GO) biological process (BP) analysis identified 13 significantly enriched summary terms, many of which were immune-related **(Figure 3C)**. Pathway analysis further revealed that the “Neuroinflammatory Signaling Pathway” and “Interferon Alpha/Beta Signaling” were activated, while pathways known to suppress microglial immunological activation, such as “IL-10 Signaling” and “PPAR Alpha-related Signaling” ^34,35^ were inhibited **(Figure 3D)**. These findings present direct evidence showing that neuroinflammation was activated in the microglia of POD patients.

**Figure 3.**
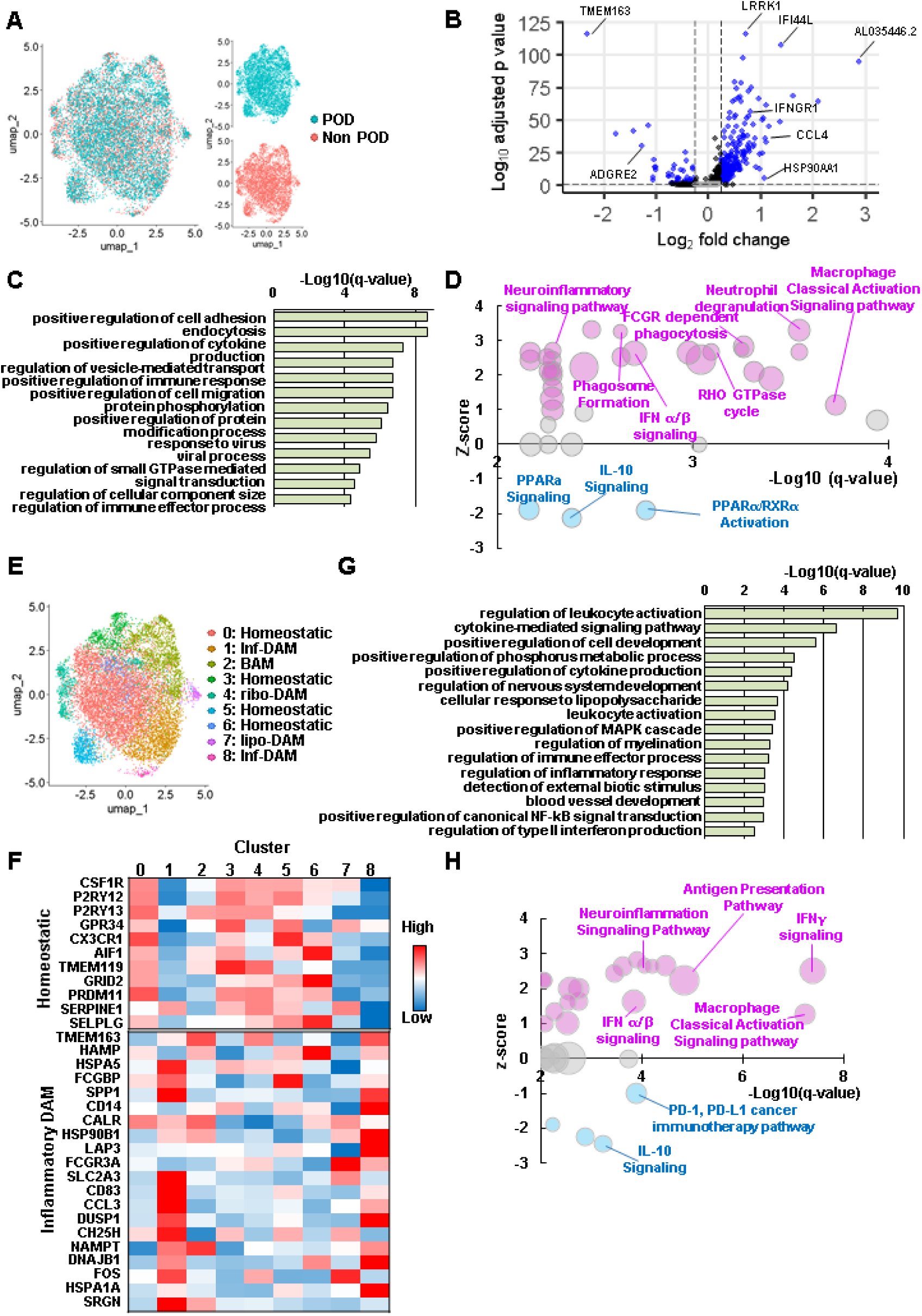
snRNAseq analysis for microglia cluster. **A** UMAP for microglia colored by the groups. **B** DEGs described in volcano plot (non POD vs POD). Blue plots meet the criteria (Log_2_FC > 0.25, adjusted p-value <0.05) **C** Enriched GO bioprocess (BP) summary terms compared between non POD and POD groups. **D** Enriched IPA pathways compared between non POD and POD groups (q-value<0.01). Bubble size means the ratio of DEGs/genes in the pathway (0.031-0.14). Magenta; activated (z-score ≥1), blue; inhibited (z-score ≤ −1), gray; random (|z-score| ≤ 1). **E** UMAP for microglia colored by clusters. **F** Relative gene expression level of representative homeostatic and inflammatory DAM markers. **G** Enriched GO bioprocess (BP) summary terms for DAM cluster. **H** Enriched IPA pathways for DAM cluster compared between non POD and POD groups (q-value<0.01). Bubble size means the ratio of DEGs/genes in the pathway (0.018 - 0.167). Magenta; activated (z-score ≥1), blue; inhibited (z-score ≤ −1), gray; random (|z-score| ≤ 1).

To determine which microglial subpopulations contributed to this transcriptional change, we performed a subcluster analysis. Recent studies have shown that disease-associated microglia (DAM) clusters emerge in both mice and humans with neuroinflammation-related neurodegenerative disorders, such as Alzheimer’s disease ^36–38^. We therefore focused on activated microglia clusters resembling DAM to investigate microglial subpopulations in greater detail. Microglia were divided into nine subclusters **(Figure 3E)**, and the distribution of cells across subclusters was similar between the POD and non POD groups **(Figure S2A)**. To characterize these subclusters, we examined the expression of representative homeostatic and inflammation-related DAM markers^38^. Clusters 1 and 8 exhibited lower expressions of homeostatic genes and higher expressions of DAM-associated genes **(Figure 3F, Figure S2B)**. We also performed trajectory analysis to examine microglial transitions among subclusters. The greatest pseudotime changes relative to Cluster 0, which was considered the major homeostatic cluster, were observed in Clusters 1 and 8 **(Figure S2C)**. Since these findings suggest that these two DAM-like clusters represent more activated and pathological microglial subpopulations, these clusters were identified as inflammatory DAM (Inf-DAM) clusters. A total of 143 DEGs were enriched in Inf-DAM clusters, with 126 genes upregulated and 17 genes downregulated in the POD group **(Figure S2D, Table S3)**. Most of the 16 terms of GO BP analysis were again immune response-related terms **(Figure 3G)**. IPA pathway analysis further confirmed that neuroinflammation was strongly upregulated in Inf-DAM microglia in the POD group **(Figure 3H)**.

Recent studies have identified additional DAM subtypes, such as ribo-DAM (ribosome-related clusters) and lipo-DAM (lysosome and lipoprotein-related clusters) in human microglia^38^. Border-associated macrophage (BAM) shares the same origin (yolk sac) and similar profiles with microglia, but are distributed in the perivascular regions and choroid plexus, distinct from parenchymal microglia ^39^. Therefore, we investigated whether the neuroinflammatory trend observed in the POD group extended to other microglial subclusters. Each subcluster was classified into five types (homeostatic, Inf-DAM, ribo-DAM, lipo-DAM, and BAM) based on the expression of representative markers **(Figure S2E, Figure 3F)**. In homeostatic microglia, enrichment analysis of 282 DEGs showed inflammatory trends such as the activation of “Neuroinflammation Signaling Pathway” and inhibition of IL-10 signaling and PPARα signaling”, which suggested a potential pre-activation state in the POD group **(Figure S2F, G, Table S4)**. This may suggest that homeostatic microglia in the POD group were stated in the pre-activation state. In the BAM-like cluster, 120 genes were enriched **(Figure S2H, Table S5)**. In the pathway analysis, since only three pathways met our original criteria (q-value ≤ 0.01), we also considered terms with a q-value below 0.05, which were deemed statistically significant. These findings suggest that neuroinflammation was also promoted in the BAM-like cluster **(Figure S2I)**. In contrast, only five genes (*TMEM163, NAMPT, SAT1, MX1, GPCPD1*) were enriched in ribo-DAM, and no genes were found to be enriched in lipo-DAM in the POD group. These findings indicate that POD-specific neuroinflammatory changes were widely occurring but unique in several subclusters to some extent.

### Transcriptional profile of astrocyte in POD

Astrocytes are also thought to play a role in the pathogenesis of delirium, as suggested by previous studies in animal models ^40,41^ and humans ^42,43^. In this study, we identified 16,080 nuclei as astrocytes based on the expression of representative markers GFAP and AQP4 **(Figure 4A**, **Figure 2C)**. Differential expression analysis revealed 734 DEGs in astrocytes from the POD group, including 536 upregulated and 198 downregulated genes **(Figure 4B, Table S6)**. GO BP analysis of astrocyte DEGs highlighted enrichment in synapse-related and morphogenesis-associated terms **(Figure 4C)**. Furthermore, IPA pathway analysis identified 52 enriched pathways, including 40 activated pathways, with synapse-related terms among the top-ranked **(Figure 4D)**. Unlike microglia, astrocytes did not show a predominant enrichment of neuroinflammation-related pathways

**Figure 4.**
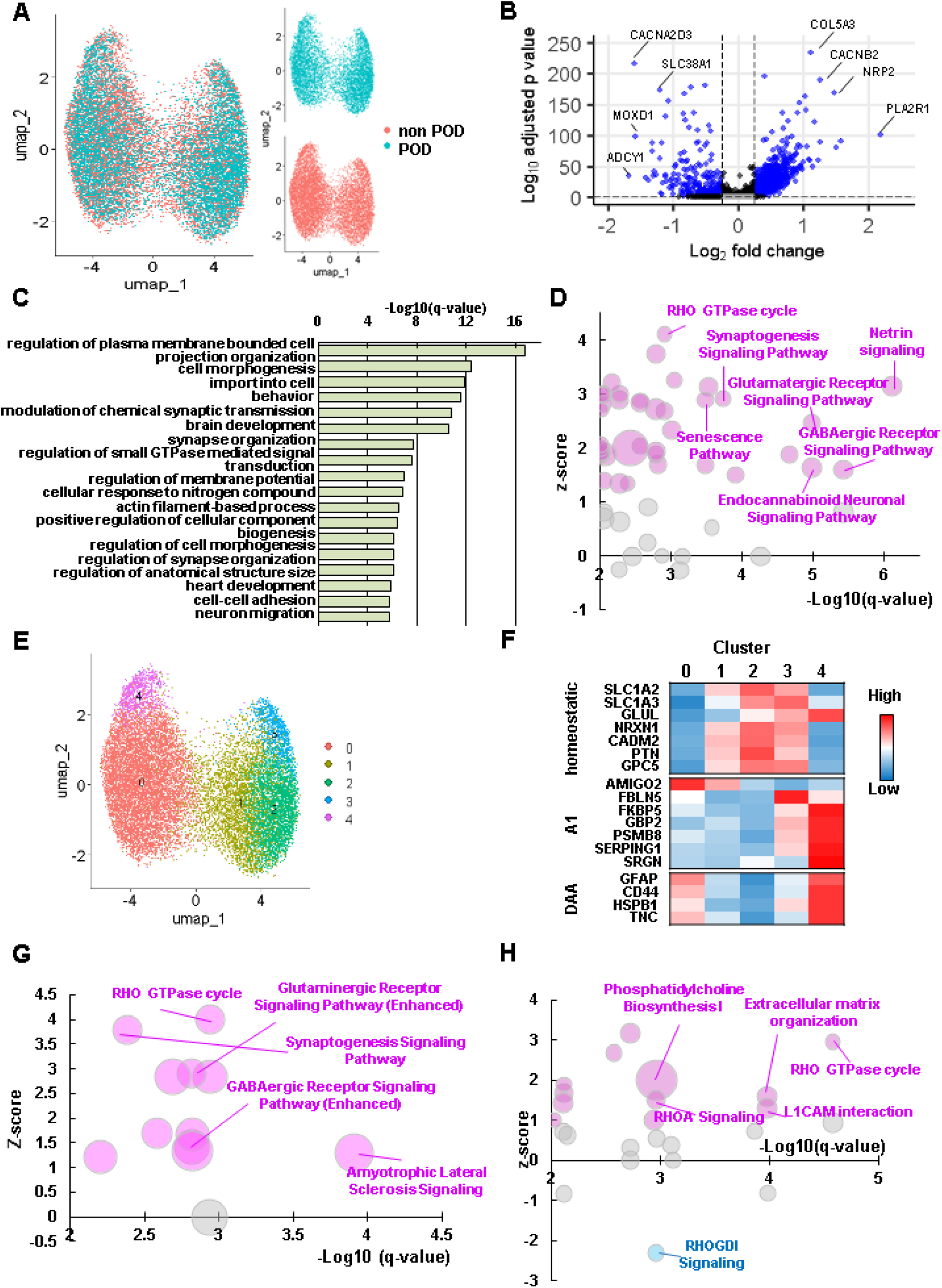
snRNAseq analysis for Astrocyte. **A** UMAP for all nuclei from astrocyte colored by groups **B** DEGs described in volcano plot. Blue plots meet the criteria (Log_2_FC > 0.25, adjusted p-value <0.05) **C** Enriched GO bioprocess (BP) summary terms compared between non POD and POD groups. **D** Enriched IPA pathways compared between non POD and POD groups (q-value<0.01). Bubble size means the ratio of DEGs/genes in the pathway (0.085-0.57). Magenta; activated (z-score ≥1), blue; inhibited (z-score ≤ −1), gray; random (|z-score| <1) **E** UMAP for all nuclei from astrocyte colored by clusters **F** Relative expression of representative markers in each astrocyte subcluster. **G** Enriched IPA pathways for homeostatic astrocytes compared between non POD and POD groups (q-value<0.01). Bubble size means the ratio of DEGs/genes in the pathway (0.0867-0.176). Magenta; activated (z-score ≥1), blue; inhibited (z-score ≤ −1), gray; random (|z-score| <1) **H** Enriched IPA pathways for activated astrocytes compared between non POD and POD groups (q-value<0.01). Bubble size means the ratio of DEGs/genes in the pathway (0.697-0.571). Magenta; activated (z-score ≥1), blue; inhibited (z-score ≤ −1), gray; random (|z-score| <1)

Reactive astrocytes, known as A1 astrocytes, are induced under neuroinflammatory conditions^44^, Recent studies have also identified disease-associated astrocytes (DAAs), which emerge in pathological states^45^. Both of these subpopulations exhibit neurotoxic phenotypes in disease contexts^46^. Additionally, activation of astrocytes has been linked to delirium-like phenotypes in a POD mouse model^40^. Given these findings, we investigated how different astrocyte subpopulations contribute to POD-related transcriptomic changes.

Astrocytes were classified into five distinct clusters **(Figure 4E)**, with a comparable distribution of cells across subclusters in both the POD and non-POD groups **(Figure S3A)**. Based on the expression of key astrocyte activation markers (A1 and DAA markers)^46^ and cluster distribution in UMAP, we characterized these subclusters as homeostatic astrocytes (clusters 1 and 2) and activated astrocytes (clusters 0 and 4) **(Figure 4F)**. In homeostatic astrocytes, we identified 1,117 DEGs **(Figure S3B, Table S7)**, which were subjected to enrichment analysis. GO BP and pathway analyses revealed significant enrichment in synapse- and neurotransmitter-related terms **(Figure S3C, Figure 4G)**. In contrast, the 734 DEGs identified in activated astrocytes **(Figure S3D, Table S8)** displayed a distinct profile, with migration-related terms being enriched and RHO GTPase signaling pathways showing characteristic upregulation **(Figure 4H, Figure S3E)**.

### Transcriptional profile of OPC and Oligodendrocyte in POD

Although the role of OPCs and oligodendrocytes in POD remains less clear compared to microglia and astrocytes, they may contribute to delirium pathogenesis through white matter dysfunction^31^. Our dataset included 8,305 nuclei classified as OPCs, characterized by the expression of the representative marker PDGFRA **(Figure S4A, Figure 2C)**. Analysis of 185 DEGs **(Figure S4B, Table S9)** revealed enrichment in several GO BP and pathway terms, including synaptic transmission-related processes **(Figure S4C, D)**. A total of 39,585 nuclei were identified as oligodendrocytes, specifically expressing MOG and MOBP **(Figure S4E, Figure 2C)**. Although 92 DEGs were detected in the POD group **(Figure S4F, Table S10)**, no significant GO or pathway terms were enriched. This suggests that, compared to other cell types, oligodendrocytes exhibit minimal transcriptional changes associated with POD.

### Downstream analysis of glial cells

According to the results of the DEG and enrichment analyses, microglia and astrocytes exhibited significant transcriptomic changes in the POD group. To assess the relationship between these changes and POD pathogenesis, we conducted disease and function enrichment analysis as a downstream analysis using the DEG lists from each cell cluster. To investigate the pathological similarities between POD and other brain-related disorders, we focused on neurological disease-associated terms that were significantly activated in the POD group (q-value < 0.01 and z-score > 1). In whole microglia, all eight enriched terms were associated with inflammation or encephalitis **(Figure 5A)**. Similarly, encephalitis-related terms were also enriched in homeostatic microglia and inflammatory DAM **(Figure 5B, C)**. Additionally, the term “cognitive impairment” was significantly enriched in homeostatic microglia, while a demyelination-related term was identified in inflammatory DAM.

**Figure 5.**
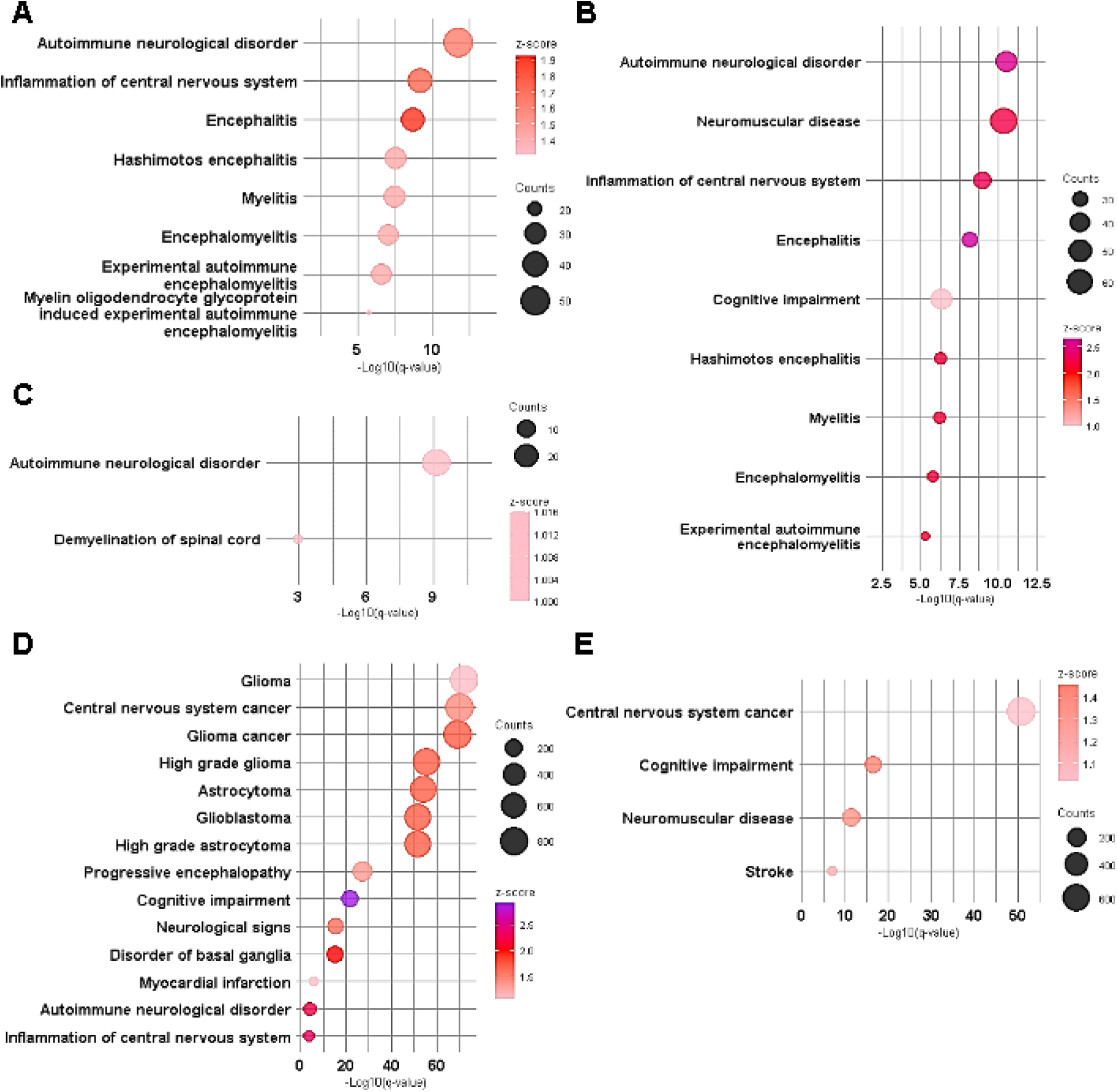
Downstream analysis for glial cells. **A-E** Enriched neurological-disease associated terms upregulated in POD group in whole microglia **(A),** homeostatic microglia **(B)**, inflammatory DAM **(C)**, whole astrocyte **(D)**, homeostatic astrocytes **(E)**. q-value <0.01. Bubble size means the gene counts of DEGs.

Enriched terms in the whole astrocyte population were primarily related to glioma and inflammation **(Figure 5D)**, whereas homeostatic astrocytes exhibited a distinct pattern **(Figure 5E)**. The only shared term between these two clusters was “Cognitive impairment.” Notably, no neurological disease-related terms were identified as upregulated in activated astrocytes.

Because delirium and dementia are often considered closely related^47^, and some glial subclusters enriched “Cognitive impairment” as expected, we further investigate the relationship between glial cells in POD and dementia, we assessed the statistical significance of five dementia-associated terms across each cell cluster **(Figure S5)**. In microglia clusters, two to three terms were significantly enriched in each cluster, suggesting that microglia in the POD group broadly exhibit transcriptional changes associated with dementia. In contrast, four to five dementia-related terms were significantly enriched in whole astrocytes and activated astrocytes, whereas none were detected in homeostatic astrocytes. This finding indicates that activated astrocytes exhibit a strong and specific transcriptional overlap with dementia pathology.

## Discussion

This study is the first to integrate DNA methylation profiling and snRNAseq in brain tissues from POD patients, providing direct molecular evidence of glial involvement in POD pathogenesis. Unlike previous studies that relied on peripheral samples or postmortem tissues, our analysis offers real-time insights from living brain tissues, allowing us to capture early molecular changes prior to delirium onset.

Our DNA methylation analysis revealed noticeable alterations in the POD group even very early on shortly after brain tissue resection and before the onset of delirium. Notably, enhanced demethylation was observed in highly methylated regions of immune-related genes, suggesting that immune response enhancement leading to POD may be regulated at the epigenetic level. Furthermore, our previous study from the same cohort has identified immune-related epigenetic changes in blood samples immediately after the brain resection surgery^9^, which were also observed in multiple other cohorts with delirium of different etiologies ^30,48^. This suggests that similar DNA methylation alterations found in the brain may be similarly detectable in the blood, raising the possibility of utilizing blood DNA methylation patterns as predictive biomarkers for delirium onset.

Our snRNAseq analysis revealed POD-specific transcriptional changes, particularly in microglia and astrocytes, whereas no significant pathway enrichment was observed in oligodendrocytes. This suggests that cell-type specificity exists in the early stages of POD pathogenesis. In microglia, we observed a strong and widespread activation of neuroinflammatory pathways. Previous studies have suggested the involvement of neuroinflammation in POD, reporting upregulated inflammatory markers such as pro-inflammatory cytokines in cerebrospinal fluid^49,50^ and postmortem brain samples^42^. While these studies indirectly linked neuroinflammation to POD, our findings provide the first direct evidence of enhanced neuroinflammatory responses in microglia during POD pathogenesis. Additionally, our enrichment analysis revealed inhibition of PPARα signaling in microglia, particularly in homeostatic microglia. PPARα signaling is known to have suppressive effects on microglial activation^51,52^. This suggests that PPARα suppression may increase microglial susceptibility to a pro-inflammatory state, thereby promoting POD pathogenesis. These findings also raise the possibility that PPARα agonists could have a preventive effect on POD.

Regarding astrocytes, multiple signaling pathways were enriched in the POD group. Astrocytes play diverse roles, including regulating synaptic activity and neurotransmission, maintaining the blood-brain barrier (BBB), mediating neuroinflammation, and energy metabolism ^31,53^. In homeostatic astrocytes, synapse-related pathways were upregulated, which may contribute to excitatory-inhibitory imbalance, a fundamental mechanism implicated in delirium^54^. Conversely, activated astrocytes exhibited RHO GTPase upregulation and enrichment of migration-related pathways. In general, RHO GTPase negatively regulates astrocyte migration^55^, whereas inflammatory stimulation is known to inhibit RHO GTPase and promote astrogliosis^56^. However, our findings showed heightened RHO GTPase activity in POD astrocytes, suggesting that RHO GTPase activity and the altered migration may suppress POD pathogenesis.

Downstream analysis revealed that POD-associated microglia and astrocytes share common transcriptional trends with several diseases, suggesting that therapeutic strategies used for related conditions may be applicable to POD treatment. The transcriptional profile of microglia in the POD group closely resembled that observed in encephalitis. While microglia are known to play both protective and detrimental roles in encephalitis, they are critical mediators of the disease process^57^. This similarity suggests that microglia may exhibit comparable behaviors in encephalitis and POD pathogenesis. Astrocytes were also implicated in inflammatory pathology, although their subclusters did not show a clear trend. Furthermore, the downstream analysis highlighted a potential link between dementia and POD. The term “cognitive impairment” was significantly enriched in both microglia and astrocyte clusters, and multiple dementia-related terms were also significantly enriched. These findings support the hypothesis of a shared pathological mechanism between dementia and POD.

This study has several limitations. First, the small sample size (18 subjects, including 9 POD patients) limits statistical power. However, analyzing brain tissue from living patients offers a unique opportunity to directly study POD pathology. To the best of our knowledge, this is the first multi-omics analysis of POD patient brain samples, marking an important step toward understanding its molecular mechanisms. Future studies with larger cohorts will be needed to validate these findings. Second, due to sample size limitations, we did not examine sex- and age-dependent differences. While age is a well-known risk factor for delirium, the influence of sex remains controversial ^58,59^. Larger studies incorporating these variables will be essential for a comprehensive understanding of POD pathogenesis. Third, no single CpG site reached genome-wide significance, likely due to the small sample size. Additionally, the challenge of detecting significant differences may stem from cell-type mixture in bulk methylation data, emphasizing the need for cell-type-specific epigenomic analyses. Nonetheless, potential CpG clusters in our analysis displayed unique characteristics related to POD, warranting further investigation. Lastly, our snRNAseq analysis focused on glial cells, leaving neurons largely unexamined. Given the nature of the resected brain tissue from epilepsy focal resection surgery, sample variability across brain regions resulted in diverse neuronal characteristics, making analysis challenging. However, since neural activity is correlated with POD^60^, targeted neuronal analysis will be crucial in future studies.

Despite these limitations, our study provides new insights into the molecular underpinnings of POD pathogenesis, highlighting the direct role of glial cells in disease development. By integrating DNA methylation and snRNAseq analysis, we uncovered direct evidence of epigenetic and transcriptomic mechanisms associated with neuroinflammation and more in POD. These findings suggest novel therapeutic strategies, particularly targeting microglia and astrocytes, to mitigate POD risk and improve patient outcomes.

## Data availability

DNA methylation data and snRNAseq data have been deposited in the Gene Expression Omnibus database (GSE290719, GSE291019). Other processed data generated in the present study are available from the corresponding author, G.S., upon reasonable request.

## Acknowledgments

The authors appreciate the patients who participated in this study. This work was supported by research grants from the National Institute of Mental Health, United States (R01 MH119165, AG084710).

---

**Figure S1.**
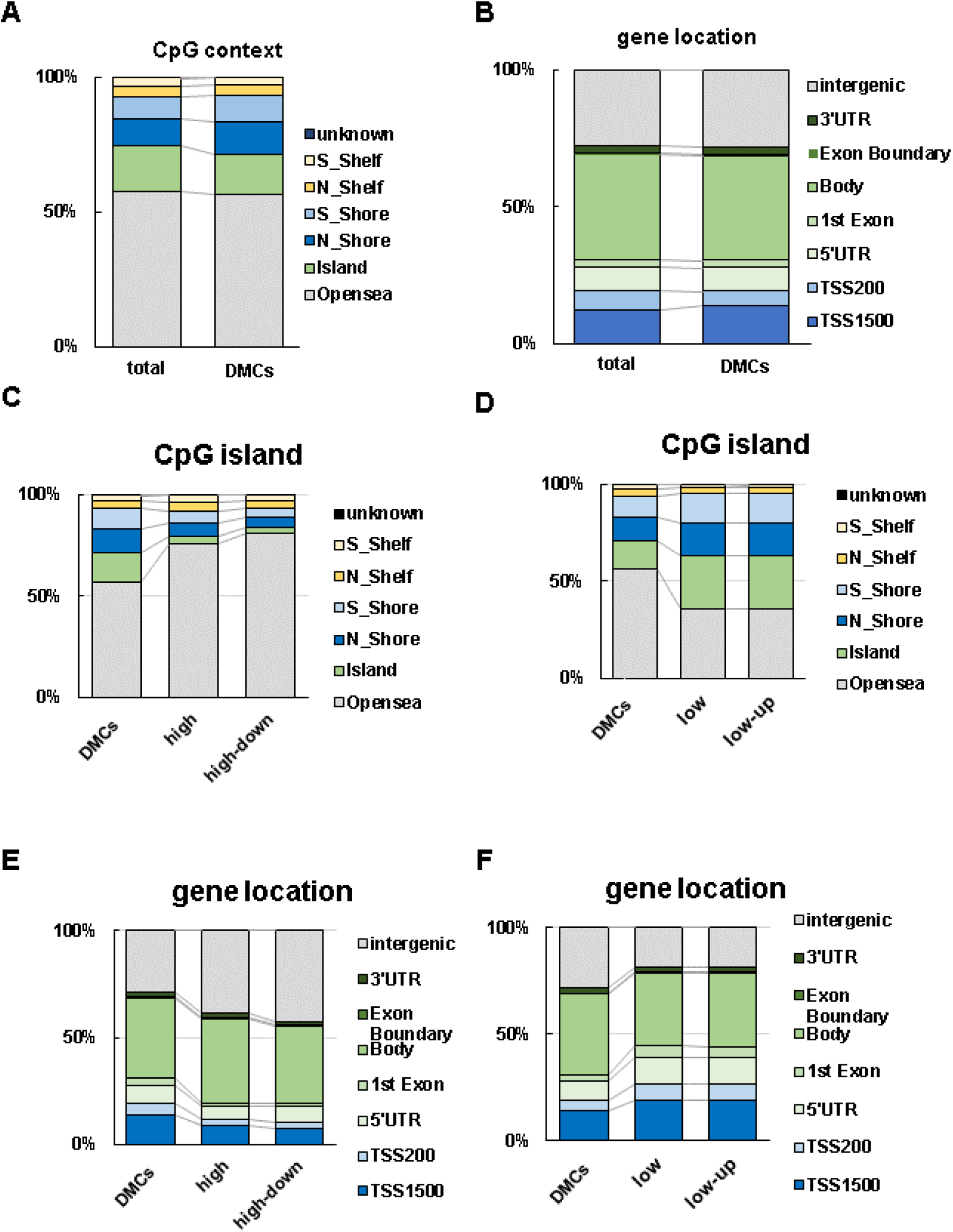
Characteristics of differentially methyrated CpGs. **A, B** the proportions of DMPs site location; A. CpG context and B. gene location. **C, D** the detailed proportion of CpG context; C.high methylated DMPs D. low methylated DMPs **E, F** the detailed proportion of gene location; E.high methylated DMPs F. low methylated DMPs

**Figure S2.**
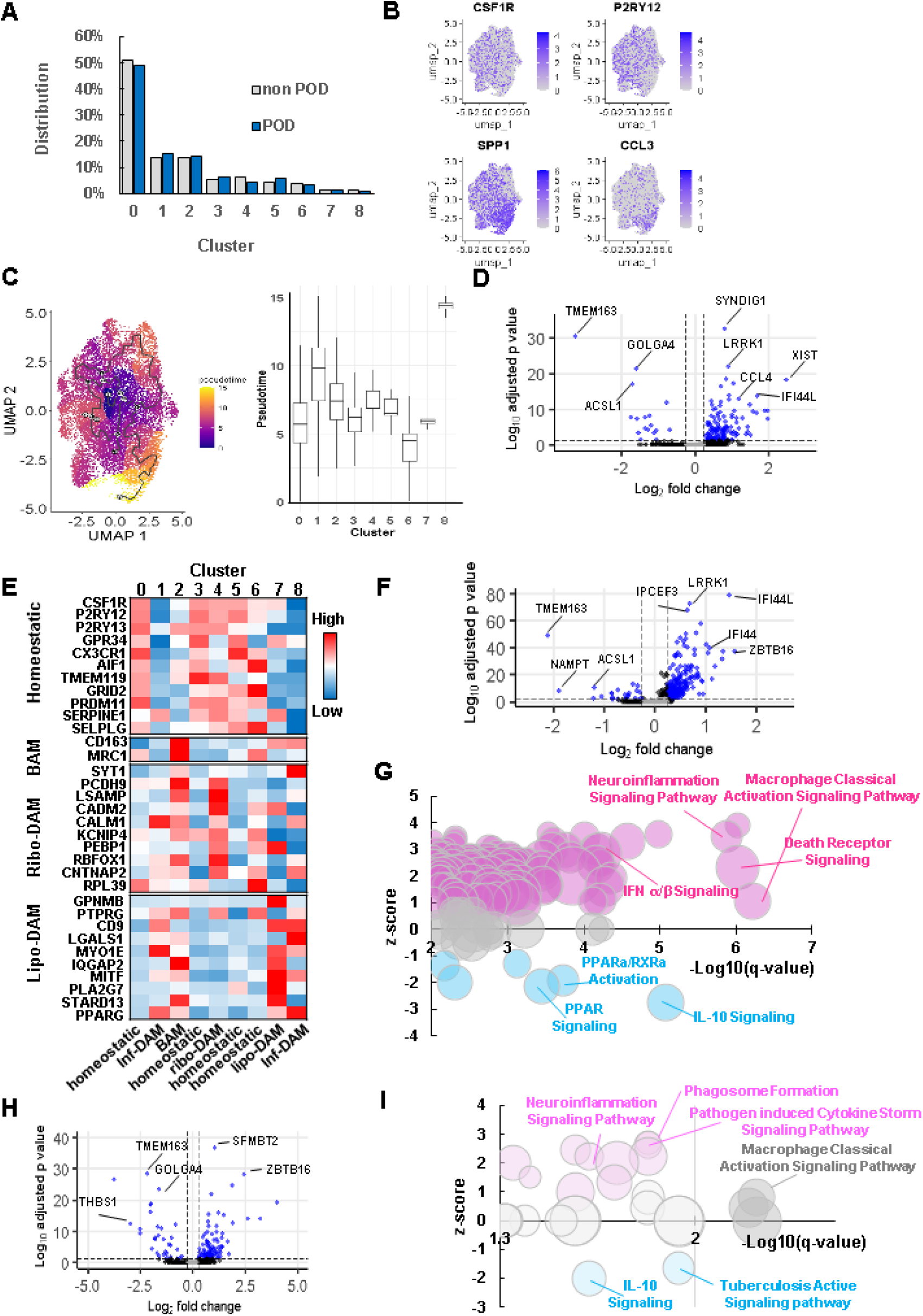
snRNAseq analysis for microglial sub-cluster. **A** Distribution of microglia in each sub-cluster **B** Representative homeostatic and DAM marker expression **C** Trajectory analysis for microglia cluster. pseudo time distribution in UMAP (left) and each cluster (right). **D** DEGs in inflammatory DAM described in volcano plot. Blue plots meet the criteria (Log_2_FC > 0.25, adjusted p-value <0.05) **E** Gene expression level of representative homeostatic, BAM and DAM markers. **F** DEGs in homeostatic microglia described in volcano plot. Blue plots meet the criteria (Log_2_FC > 0.25, adjusted p-value <0.05) **G** Enriched IPA pathways in homeostatic microglia cluster (q-value<0.01). Bubble size means the ratio of DEGs/genes in the pathway (0.0271-0.208). Magenta; activated (z-score ≥1), blue; inhibited (z-score ≤ ࢤ1), gray; random (|z-score| ≤ 1). **H** DEGs in BAM described in volcano plot. Blue plots meet the criteria (Log_2_FC > 0.25, adjusted p-value <0.05) **I** Enriched IPA pathways in BAM cluster compared between non POD and POD groups (q-value<0.01). Bubble size means the ratio of DEGs/genes in the pathway (0.0128-0.0727). Magenta; activated (z-score ≥1), blue; inhibited (z-score ≤ −1), gray; random (|z-score| ≤ 1).

**Figure S3.**
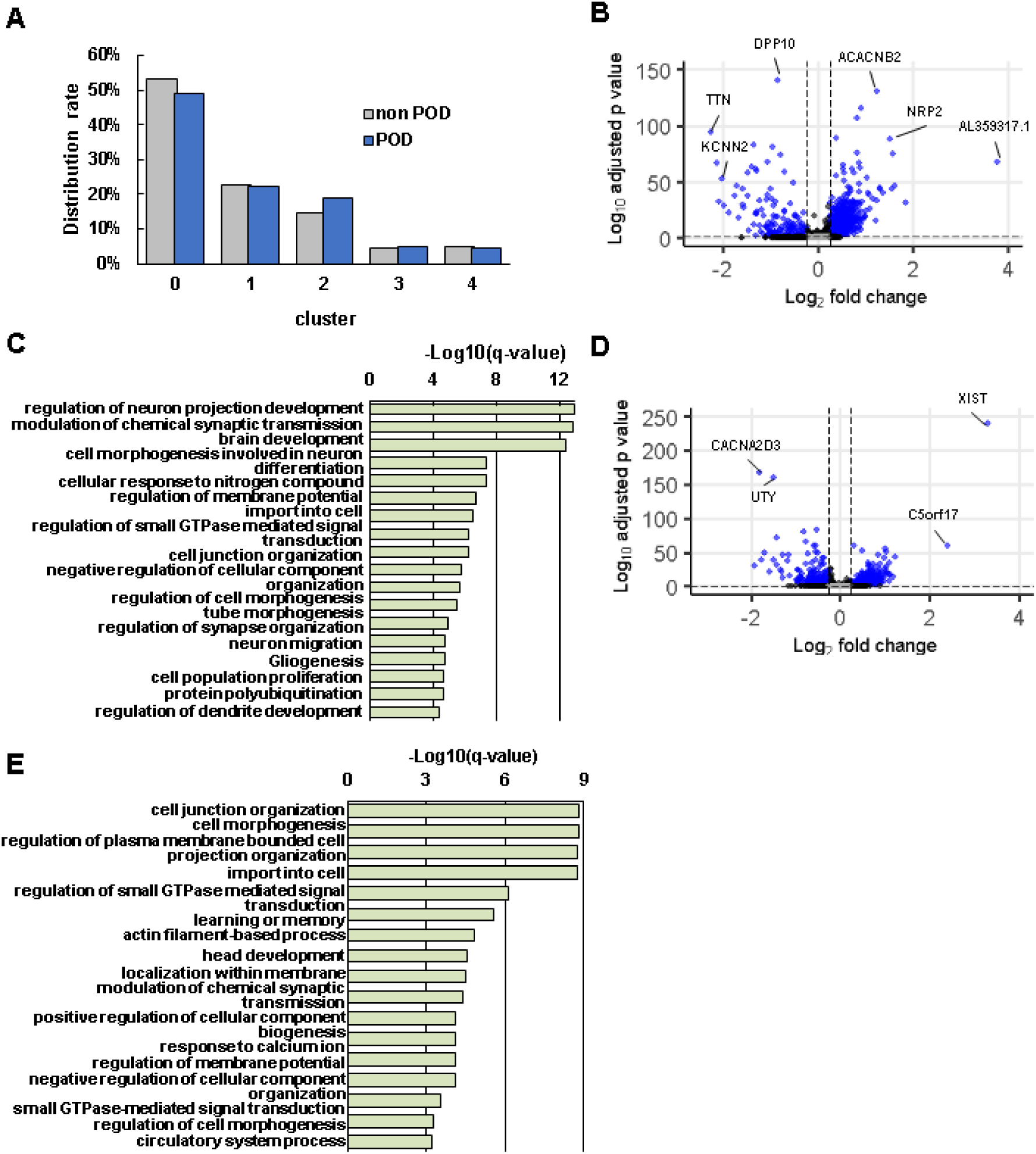
snRNAseq analysis for astrocyte sub-cluster. **A** Distribution of astrocyte in each sub-cluster **B** DEGs in basic astrocyte cluster described in volcano plot. Blue plots meet the criteria (Log_2_FC > 0.25, adjusted p-value <0.05) **C** Enriched GO bioprocess (BP) summary terms in homeostatic astrocytes **D** DEGs in activated astrocyte cluster described in volcano plot. Blue plots meet the criteria (Log_2_FC > 0.25, adjusted p-value <0.05) **E** Enriched GO bioprocess (BP) summary terms in activated astrocytes **F** Relative expression of representative markers in each astrocyte subcluster

**Figure S4.**
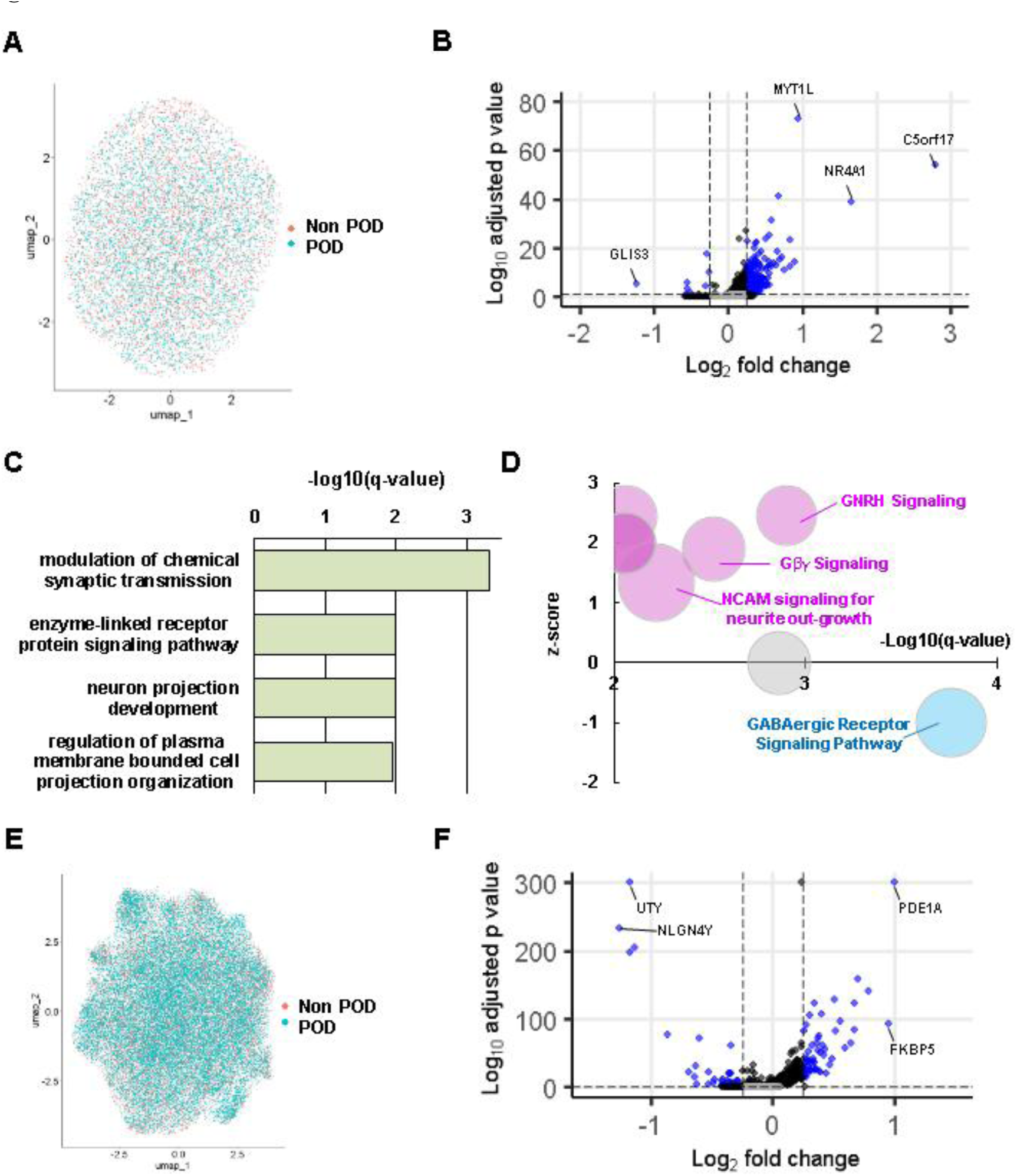
snRNAseq analysis for OPC and Oligodendrocyte cluster. **A** UMAP for all nuclei from OPCs colored by groups **B** DEGs in OPCs described in volcano plot. Blue plots meet the criteria (Log_2_FC > 0.25, adjusted p-value <0.05) **C** Enriched GO bioprocess (BP) summary terms in OPCs **D** Enriched IPA pathways in OPCs (q-value<0.01). Bubble size means the ratio of DEGs/genes in the pathway (0.0412-0.0794). Magenta; activated (z-score ≥1), blue; inhibited (z-score ≤ −1), gray; random (|z-score| < 1) **E** UMAP for all nuclei from oligodendrocyte colored by groups **F** DEGs described in volcano plot. Blue plots meet the criteria (Log_2_FC > 0.25, adjusted p-value <0.05)

**Figure S5.**
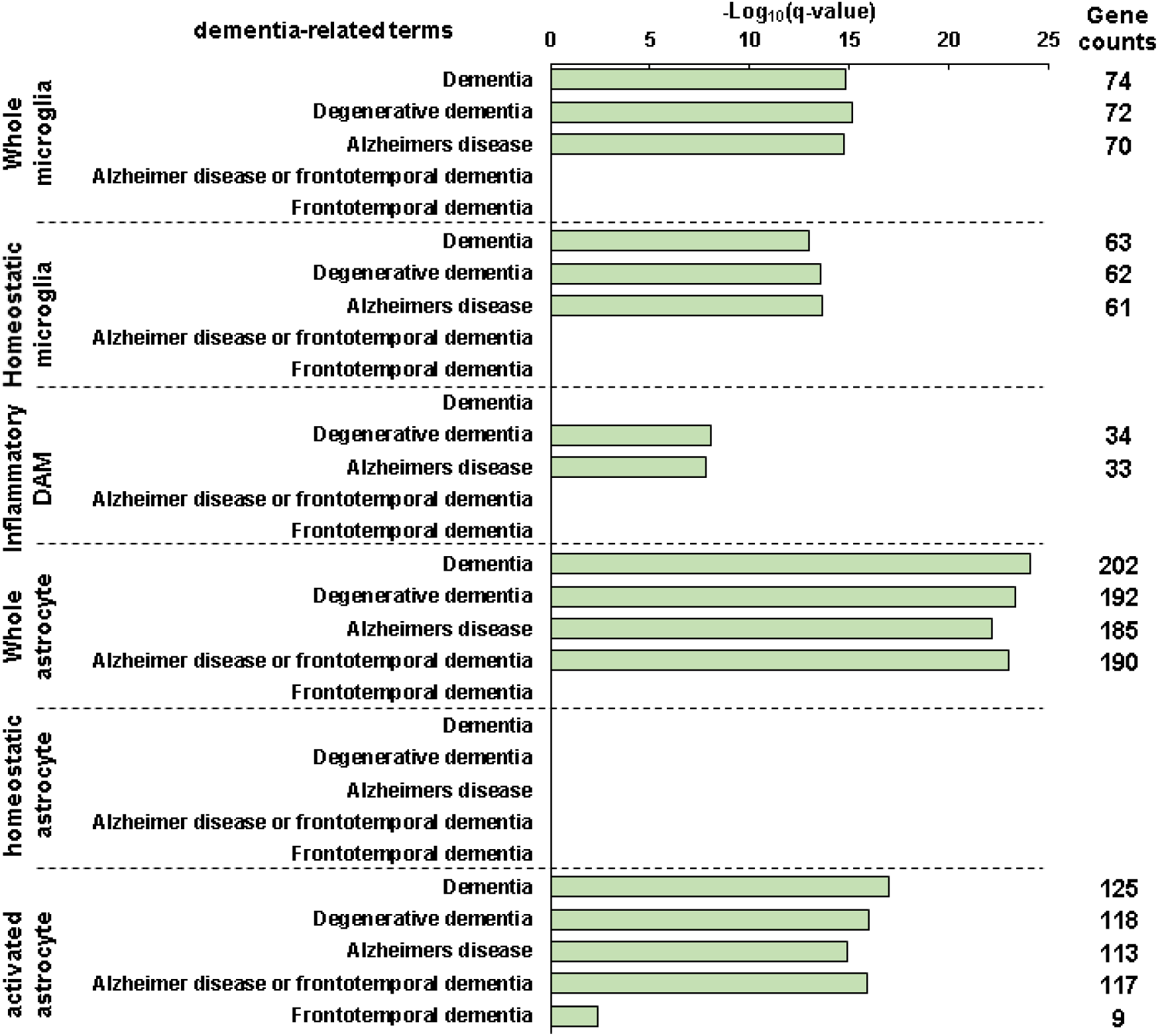
Dementia-related terms in microglia and astrocytes. Statistical significance of 5 dementia-related terms in microglia and astrocytes clusters.

